# CARD k-mers: Unmasking the pathogen hosts and genomic contexts of antimicrobial resistance genes in metagenomic sequences

**DOI:** 10.1101/2025.09.15.676352

**Authors:** Mateusz A. Wlodarski, Tammy T. Y. Lau, Brian P. Alcock, Amogelang R. Raphenya, Tiffany E. Ta, Finlay Maguire, Robert G. Beiko, Andrew G. McArthur

## Abstract

Antimicrobial resistance (AMR) is a global health crisis requiring rapid surveillance across human, agricultural, and environmental systems. A major challenge during outbreaks is not only detecting antimicrobial resistance genes (ARGs), but also unmasking their pathogen hosts and genomic context, as ARGs alone do not fully capture AMR risk. Pathogen identification is often essential for guiding effective treatment. While culture-based methods remain the diagnostic gold standard, they are slow and sometimes impractical. Faster metagenomic (mNGS) tools typically detect either ARGs, taxonomy, or genomic context, but rarely all three, resulting in fragmented surveillance. Existing k-mer classifiers like Kraken2 and CLARK, designed for general taxonomy, often perform poorly on AMR-specific sequences. We introduce CARD k-mers, the first tool built to jointly predict species-level taxonomy and genomic context (plasmid vs. chromosome) for ARGs in short metagenomic reads. Integrated with the Comprehensive Antibiotic Resistance Database (CARD), CARD k-mers enables rapid, context-aware assignment of ARGs to their likely pathogen and genomic element origin. In benchmarking with 103,456 *in-*silico pathogen-specific AMR alleles, CARD k-mers outperformed Kraken2 and CLARK by 10.85% and 15.2%, respectively, and correctly classified the genomic context of 4,590 chromosome- and 176 plasmid-specific ARGs. The tool operates at speeds exceeding 675,000 metagenomic reads per minute. By delivering fast, accurate, and context-rich classification of ARGs, CARD k-mers significantly advances untargeted AMR surveillance and is accessible to users with basic command-line experience for use in both clinical and environmental pipelines. CARD k-mers is available at: https://github.com/arpcard/rgi.

## INTRODUCTION

AMR is a pressing global health challenge exacerbated by the overuse of antibiotics and a stagnation in the development of novel drugs (1, 2). Without decisive mitigation efforts, AMR is projected to cause approximately 39 million cumulative deaths by 2050 (1). Antibiotic-resistant bacterial pathogens harboring ARGs evade treatment through well-characterized resistance mechanisms (3), with particular concern for plasmid-borne elements due to their high potential mobility between organisms via lateral gene transfer (LGT) (4). These events facilitate the rapid dissemination of resistance, leading to multidrug-(MDRO) and ultimately, to extensively drug-resistant (XDRO) organisms (5).

Recent advances in sequencing technologies now enable comprehensive genomic surveillance of microbial environments, from wastewater and agricultural runoff to clinical samples (6, 7). These methods facilitate the detection and tracking of ARGs and priority AMR pathogens across diverse contexts. Resources such as the Comprehensive Antibiotic Resistance Database (CARD) have become indispensable in this effort, offering tools like the Resistance Gene Identifier (RGI, CARD’s main software) for *in-silico* identification of known ARGs and putative novel variant discovery (8). While these advancements have aided global public efforts, a critical gap remains linking detected ARGs to their pathogen hosts and genomic contexts directly from metagenomic data. This step is essential in clinical settings, where timely and accurate diagnoses are necessary to guide effective antibiotic treatment (6).

Historically, pathogen and plasmid identification relied on cultured isolates, where growth media and molecular diagnostics inherently provided taxonomic information and genomic context (9). These traditional phenotypic methods are limited by their reliance on culturable bacteria while being labor-intensive (10). Consequently, culture-independent, genotypic methods are increasingly replacing traditional approaches due to their ability to sequence entire microbial communities with high-throughput and speed (2). These mNGS datasets, often containing millions of sequencing reads, present unique analytical challenges for determining the pathogen hosts and genomic contexts of ARGs.

For decades, alignment-based tools such as BLAST, Burrows-Wheeler Transform, and Hidden Markov Model searches have been the dominant approaches for sequence classification (11). Alignment methods rely on aligning DNA reads to a reference genome to assign taxonomic labels. However, they are computationally intensive, rely on subjective scoring matrices, and assume homology between sequences, making them ill-suited for analyzing large, diverse metagenomes (6, 11, 12). LGT events in microbial populations often violate core assumptions of alignment-based approaches, further limiting their utility (11).

To address these challenges, bioinformatics has shifted toward alignment-free sequence classification methods, particularly k-mer-based approaches, also used in metagenomic assembly and a genome-wide association studies (13–15). These methods rely on exact word matching between pre-computed pathogen-specific k-mers and query sequences, bypassing the computational bottlenecks of alignment tools. Kraken2 is a leading k-mer-based classifier, capable of processing millions of reads per minute, making it highly suitable for large-scale sequence analysis (16, 17). Kraken2 classifies sequences using the lowest common ancestor approach, assigning reads to the lowest common taxonomic rank shared by all organisms containing a given k-mer. Conversely, CLARK, an alternative k-mer classifier, relies on discriminatory k-mers, which are uniquely associated with specific taxa. However, recent evaluations have demonstrated that k-mer tools rely heavily on general-purpose reference databases that are not optimized for domain-specific analyses (18–20). This dependency can severely limit their accuracy and precision when applied to AMR-focused datasets, where the taxonomic and functional complexity often surpasses what broad-spectrum reference libraries can effectively capture. This highlights the need for a specialized classifier optimized for ARGs and their associated pathogens, enabling untargeted AMR surveillance of metagenomic samples.

CARD’s RGI software now supports a k-mer classification algorithm (CARD k-mers) specifically tailored for metagenomic AMR surveillance. CARD k-mers integrates seamlessly with CARD’s Resistomes & Variants sequence data (CARD-R, https://card.mcmaster.ca/prevalence), enabling accurate species-level taxonomic predictions and mobility analysis for metagenomic reads encoding ARGs. CARD-R reflects computer-generated resistome predictions for 414 important pathogens and includes ARG sequence variants beyond those reported in the scientific literature. By leveraging CARD-R, ARG alleles matching the RGI-Strict criteria, representing ARG variants closely related to known alleles but not yet described in the literature, CARD k-mers is uniquely positioned to detect pathogens with both known and novel functional variant ARGs.

CARD k-mers operates by constructing a reference k-mer set derived from the alleles stored in the CARD-R database, a massive pathogen-centric database containing predicted ARG alleles. For CARD-R version 4.0.0, this dataset includes over 40 million unique k-mers of length 61 (61-mers, Supplementary Table 1). These k-mers are divided into two primary categories (taxonomic versus genomic k-mers), with each serving a specific role in classification. Taxonomic k-mers are designed to predict the pathogen-of-origin and consist of single-species k-mers, which are unique to ARG alleles from individual species, and genus-specific k-mers, which are unique to ARG alleles from a specific genus. These classifications allow for both species-level identification and broader taxonomic resolution when species-specific resolution is not possible. Genomic k-mers, on the other hand, focus on the genomic context of ARG alleles and their potential mobility. They are further categorized into k-mers unique to chromosomally-encoded ARG alleles, k-mers unique to plasmid-encoded ARG alleles, and k-mers shared between chromosome- and plasmid-encoded ARG alleles. ARGs that display little to no allelic variation and that are widely distributed across diverse taxa, such as NDM-1, are not suitable for pathogen-of-origin prediction but can be assessed for genomic context to generate mobility risk insights. It is important to emphasize that both taxonomic and genomic k-mers are derived from ARG allele sequences only and not any flanking sequences, reserving analysis and predictive ability to the specific domain of ARG alleles. This dual classification strategy enables CARD k-mers not only to identify the pathogen-of-origin but also to evaluate the mobility risks of predicted ARG alleles.

## MATERIALS AND METHODS

### RGI Software and Reference K-mers

The CARD k-mers algorithm is implemented as a command-line tool within CARD’s RGI software (*rgi kmer_query*) and depends on reference k-mers derived from the CARD-R database, as outlined above. Users can download the CARD-R data from the CARD website and configure custom reference k-mers of any desired length using the command *rgi kmer_build*, but the CARD website provides pre-computed 61-mers for download for each CARD-R update. The validation described in this manuscript uses these pre-computed 61-mers except where specified.

User input sequences are classified using CARD k-mers to infer the most likely pathogen host and determine whether each AMR gene is chromosome- or plasmid-borne (Figure 1). Classification decisions follow a structured rules-based logic tree (Supplementary Figure 1). The *rgi kmer_query* algorithm accepts input files in multiple formats, including FASTA, RGI JSON, and RGI bwt BAM, and produces output files in both tab-delimited text and JSON formats (Figure 2). These outputs provide a summary of the pathogen-of-origin predictions and k-mer counts for chromosomal and plasmid sequences associated with the input data. For taxonomic predictions, input sequences must meet a minimum k-mer coverage threshold (default of 10 k-mers; user adjustable).

**Figure 1.**
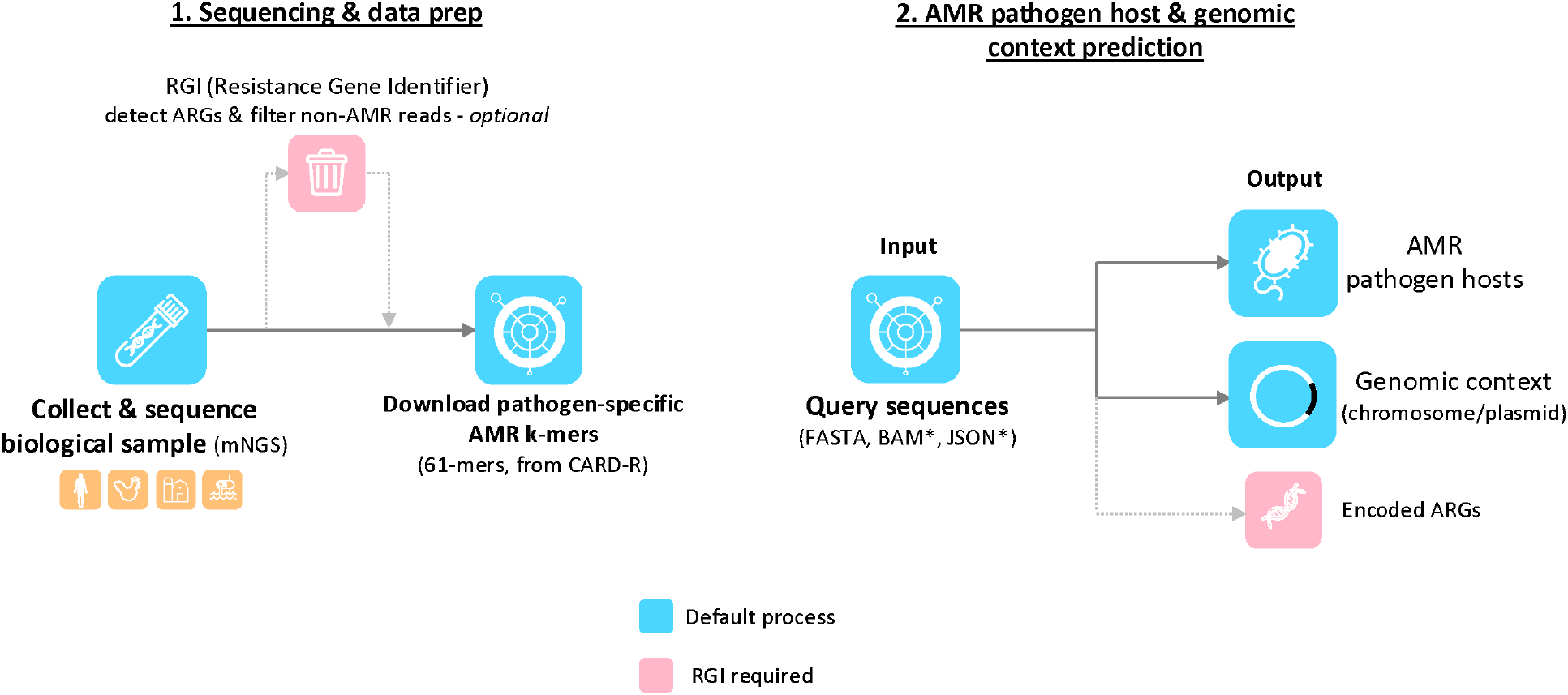
Overview of a metagenomic AMR surveillance workflow using CARD k-mers for pathogen and genomic context prediction. Step 1: A biological sample is collected and subjected to metagenomic sequencing (mNGS). Recommended but optional, the Resistance Gene Identifier (RGI) can be used to detect AMR genes and filter out non-AMR reads prior to analysis. Pathogen-specific AMR k-mers (61-mers from CARD-R) are downloaded to construct the reference database. Step 2: Sequences are queried using CARD k-mers, supporting input formats such as FASTA, BAM, or JSON. The tool predicts both the pathogen host and the genomic context (chromosome vs. plasmid) of the AMR genes. Encoded ARGs can also be inferred if RGI filtering is applied upstream.

**Figure 2.**
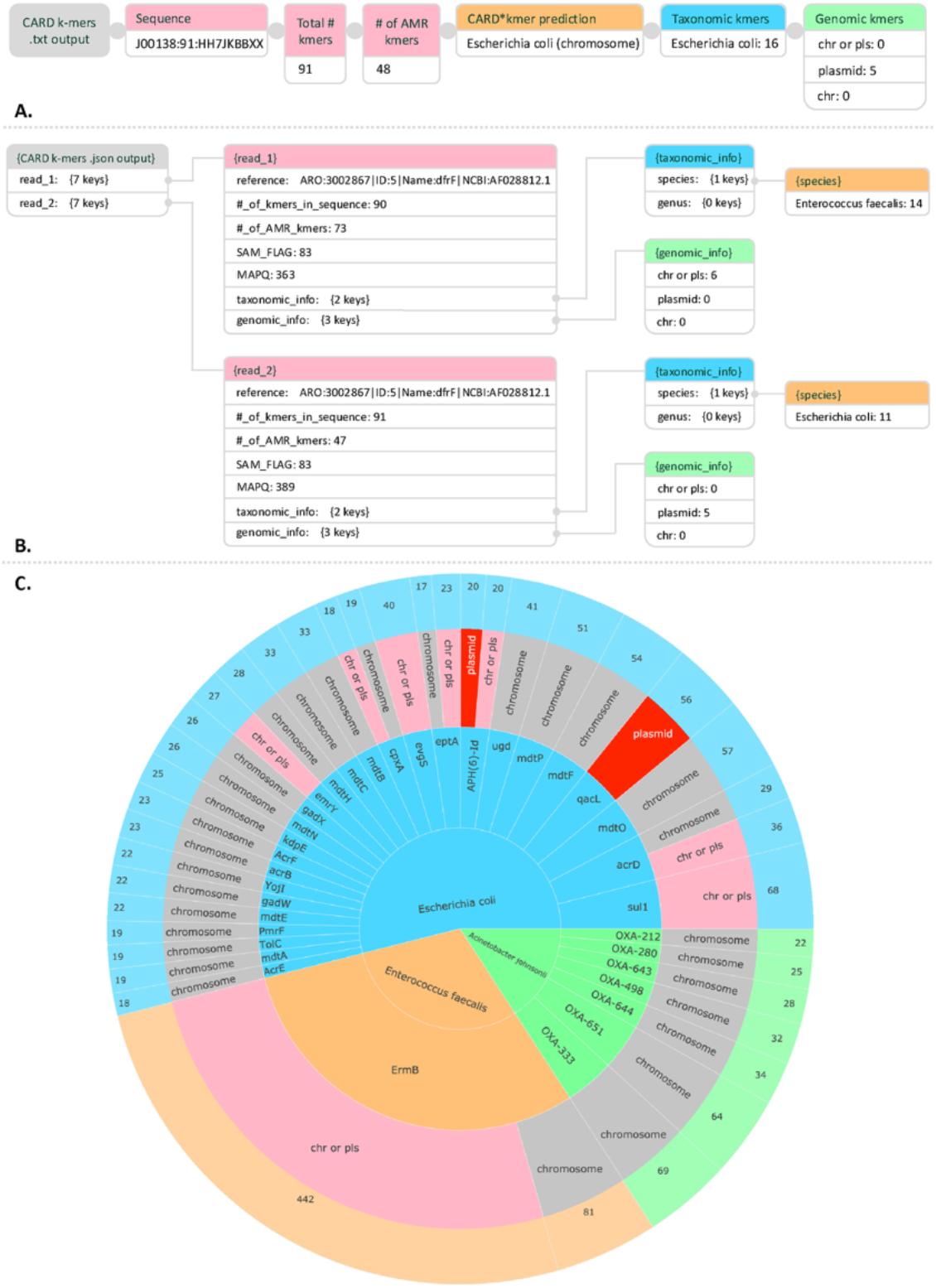
CARD k-mers multi-format outputs and visualization potential. (A) Structure of text output from the CARD k-mers tool, summarizing per-sequence information such as total k-mers, AMR-specific k-mers, CARD-based predictions, taxonomic assignments (e.g., *Escherichia coli*), and genomic context (e.g., chromosome or plasmid). (B) Structure of JSON output displaying detailed k-mer counts per species, genus, and genomic context. Unlike the text summary, this format provides a granular view of the k-mer evidence across possible origins, without making direct classification calls. (C) Example sunburst chart generated from mock CARD k-mers results on metagenomic data, illustrating the tool’s capacity to support downstream visualizations. Concentric layers represent the detected AMR gene (center), associated pathogen, predicted genomic context (chromosome or plasmid), and read-level contribution. This preview from the (in-development) *k-mer viz* module demonstrates how users will be able to explore resistance patterns, taxonomic distributions, and genomic origins interactively.

### Classification Validation Datasets

Two datasets were constructed to perform separate validation tests for CARD k-mers: pathogen and genomic classification. The first dataset to validate pathogen classification accuracy included 320,614 *in silico* predicted ARG alleles, sourced from CARD-R version 4.0.0 (Figure 3). As an initial filter, 9,722 alleles were excluded because of probable LGT occurrence, since they were observed in multiple species, making them non-pathogen specific. For example, allele “A” was excluded as it appeared in 48 different pathogens, whereas allele “B”, unique to *Salmonella enterica*, was retained. A random two thirds of the sampled alleles (rounded down) for each pathogen were allocated to a “reference set” to build a 61-mer reference library of taxonomic and genomic k-mers using *rgi kmer_build*. The remaining one-third was assigned to a “testing set” for validation. To ensure inclusion, each pathogen required at least three unique alleles in CARD-R, the minimum necessary for our one-third holdout validation strategy. As a result, 62 pathogens were excluded due to insufficient data, leaving a total of 315 pathogens in the validation dataset. Stratifying the data by pathogen preserved the relative allele distribution rates of the original CARD-R dataset. Altogether, the training set contained 207,589 alleles from 315 pathogens, while the testing set included 103,456 alleles to validate classification accuracy. For taxonomic validation, RGI v6.0.2 *kmer_query* was compared against two Kraken2 instances and one CLARK instance. Kraken2 default utilized Kraken2’s standard 70 GB library (version 2.0.8) while Kraken2 (CARD) was constructed from the training set containing 207,589 ARG alleles as outlined above. CLARK used its standard library (version 1.3.0). For genomic classification accuracy, a subset of 8,207 genomic-specific alleles (i.e., alleles found exclusively in chromosomes or plasmids) from the testing set were used (Figure 3). As the original CARD-R data included pathogen, chromosome, and plasmid prevalence for each ARG allele sequence, accuracy was defined as *rgi kmer_query* predicting the same taxonomic or genomic classification while inaccuracy was defined as disagreement between CARD-R and *rgi kmer_query*, albeit the latter can include “no classification” as possible output, which we tracked separately.

**Figure 3.**
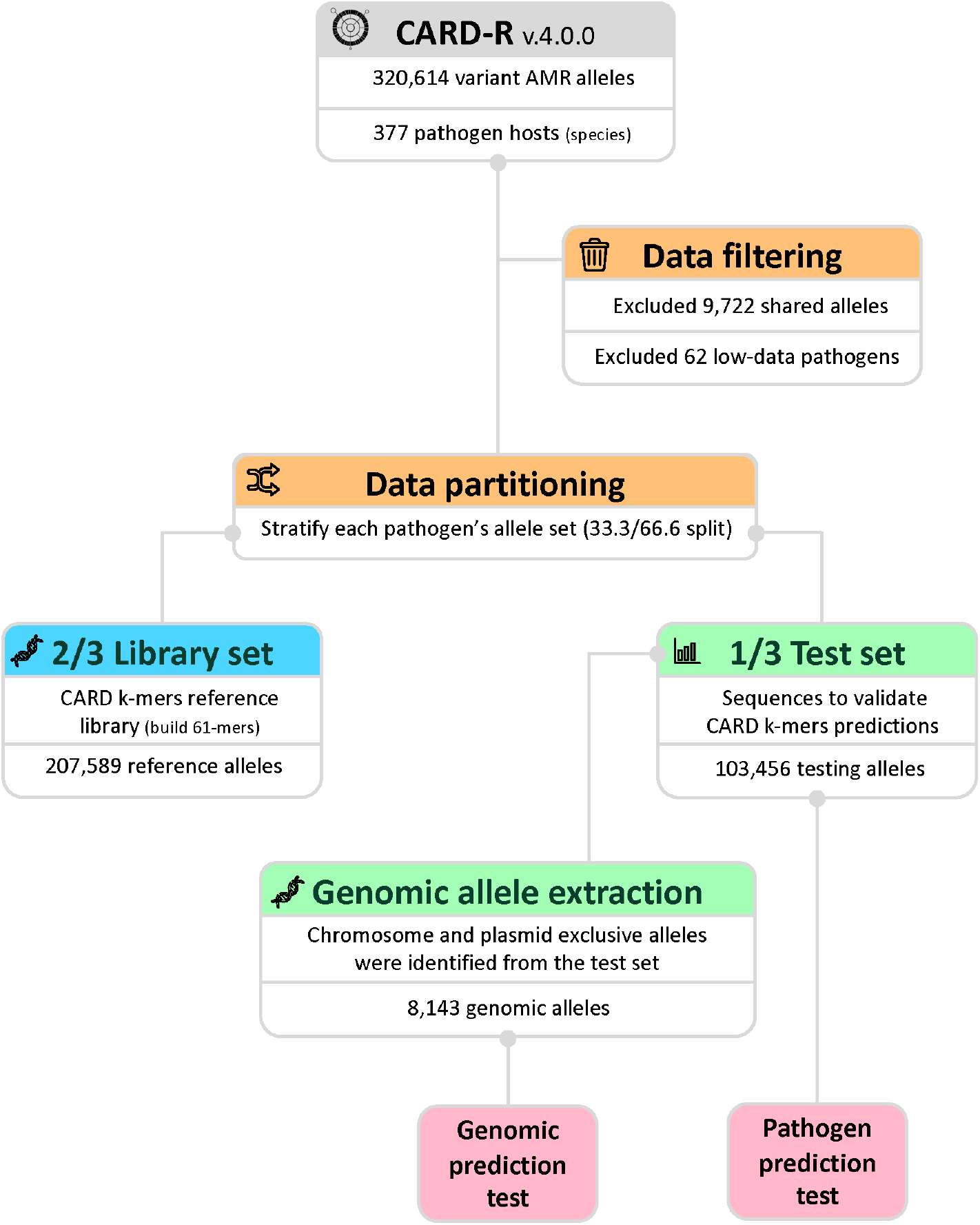
Construction of CARD k-mers validation datasets. Starting from CARD-R v4.0.0, which includes 320,614 variant AMR alleles across 377 pathogens, 9,722 shared (multi-pathogen) alleles and 62 low-data pathogens were excluded. The remaining alleles were stratified by pathogen using a by-thirds partitioning strategy, with 2/3 assigned to a reference library and 1/3 to a testing set. The resulting training set (207,589 alleles) formed the CARD k-mers reference library. The testing set (103,456 alleles) was used to validate pathogen-of-origin predictions. From this same test set, 8,143 chromosomal- or plasmid-specific alleles were extracted for evaluation of genomic origin prediction accuracy.

### Classification Rules

Following the classification of the 103,456 test set alleles, output files containing pathogen host predictions were generated by each of the four classifiers: CARD k-mers, Kraken2, Kraken2 (CARD), and CLARK. Kraken2 classifiers assigned highly specific taxonomy labels, often down to the exact strain. To standardize comparisons, an intermediate taxonomy mapping was created using ENTREZ and NCBI’s taxonomy database to scrape taxonomy identifiers and corresponding scientific names for Kraken2 and CLARK.

Classification accuracy was assessed by comparing the predicted pathogen host to the labelled pathogen in CARD-R for each allele. For Kraken2 classifiers, a species-level prediction was considered correct if there was an exact scientific name match. Since alleles in CARD-R are mapped to a base species, Kraken2 predictions specifying a strain name were also accepted as correct if the base species matched. For example, if Kraken2 predicted the pathogen of an allele as *Escherichia coli s8 0145:H28 str. RM12581*, this was considered a correct pathogen prediction because the CARD-R label for this allele was *Escherichia coli*.

Genus-level classification accuracy was recorded using similar criteria, requiring an exact genus match between the predicted pathogen and the CARD-R label genus. False species predictions were labelled as erroneous calls if an incorrect species and genus name was produced, labelled unclassified if the prediction was ambiguous, and rejected (CARD k-mers only) if the query sequence failed to meet the minimum coverage threshold value.

For the genomic classification accuracy test, accuracy was evaluated based on the ability of CARD k-mers to correctly classify the genomic context of each allele as either plasmid or chromosome. Predictions of “chromosome & plasmid” indicated that the algorithm found genomic k-mers within input sequences belonging both to plasmids and chromosomes, whereas “no genomic info” predictions were deemed as uninformative.

### Performance Benchmarking

Reference k-mer libraries from CARD-R version 4.0.0 were constructed using 61-mers and 15-mers to benchmark speed and peak memory usage during library construction.

Additionally, one million 250 base pair reads from an existing benchmarking data set (https://zenodo.org/records/6543357) (21) were classified to assess speed and peak memory usage during query processing. All benchmarking analyses were performed on a Cisco Blade server (Intel(R) Xeon(R) Gold 6238 CPU @ 2.10GHz, 88 cores, 1.5 TB RAM, running Ubuntu).

### Optimal k-mer Size

To experimentally determine optimal k-mer size (or at minimum justify the selection of 61-mers), pathogen classification validation was repeated with k values ranging from 5 to 100. An additional analysis to determine the theoretical minimum k-mer size for pathogen-specific alleles used cumulative relative entropy (CRE) analysis across millions of resampled alleles, applying a CRE cutoff of 0.1 to identify optimal thresholds using KITSUNE (22, 23).

## RESULTS

### Classification Accuracy

We evaluated the accuracy of CARD k-mers, Kraken2 (default and CARD references), and CLARK across five metrics for pathogen classification: correct species accuracy, correct genus accuracy (alleles with a correct genus, but false species prediction), erroneous predictions (wrong species and genus), unclassified alleles, and rejected sequences (applicable only to CARD k-mers) (Figure 4). CARD k-mers classified 75.69% of ARG alleles to the correct species, with an additional 3.0% to the correct genus level, and achieved the lowest error rate (1.3%) among all tools. Kraken2 (CARD) had a slightly higher species-level accuracy (78.85%) but also a higher error rate (18.72%). Kraken2 default and CLARK had lower species-level accuracies (64.84% and 60.5%, respectively) and higher erroneous predictions (31.75% and 2.0%, respectively). CARD k-mers and CLARK had the highest unclassified rates (19.88% and 34.1%, respectively), while Kraken2 tools returned fewer unclassified sequences (1.60% default and 0.11% CARD, respectively).

**Figure 4.**
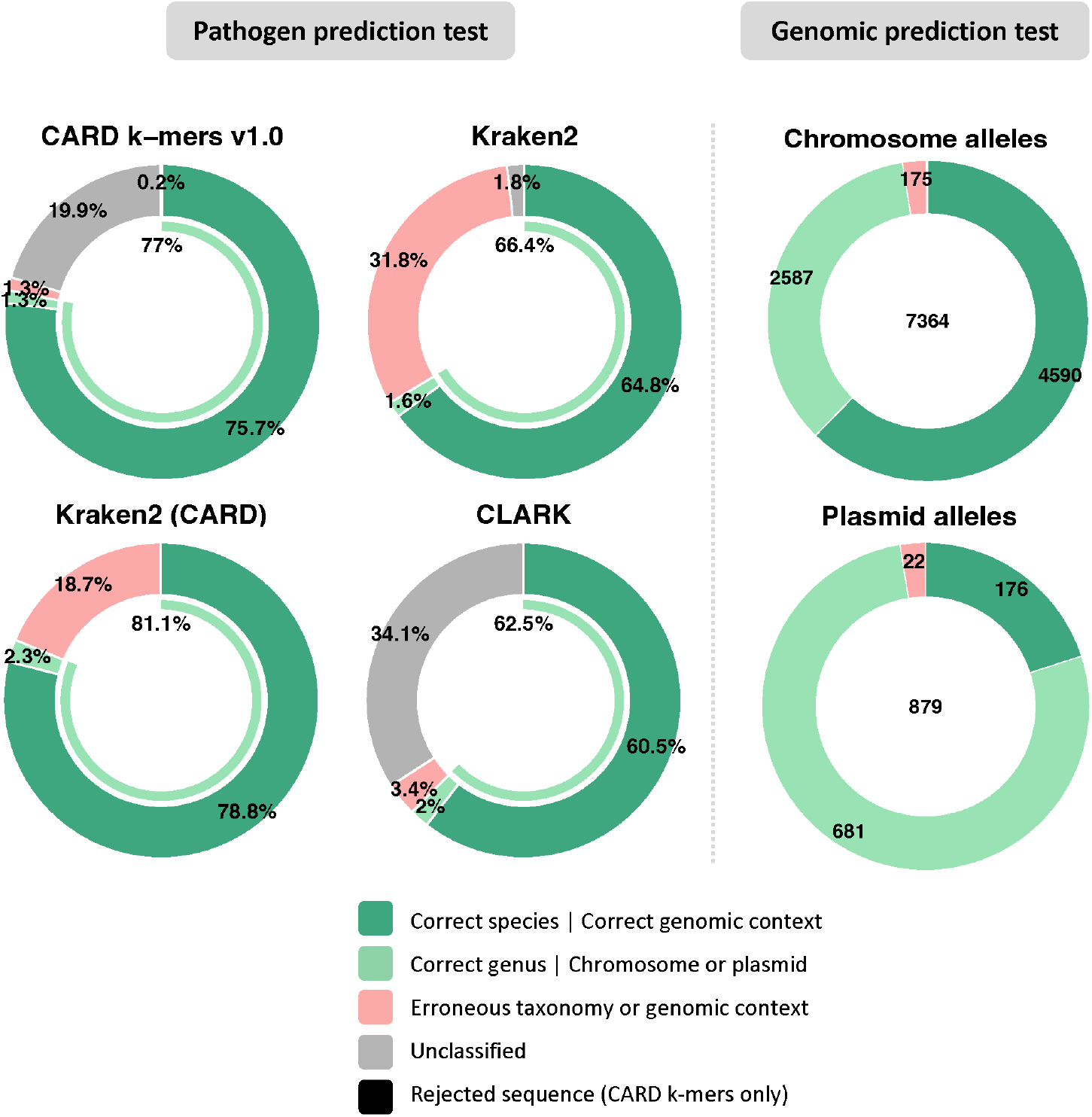
Accuracy of pathogen and genomic context classification. Classification performance is shown for pathogen prediction (left) and genomic context prediction (right) across four classifiers: CARD k-mers (61-mers), Kraken2 (with default or CARD references; both 35-mers), and CLARK (31-mers). For pathogen prediction, alleles were classified as rejected (CARD k-mers only, due to insufficient k-mers), unclassified, erroneous (incorrect species and genus), genus-level only (correct genus, incorrect species), or correct species with correct genomic context. The inner arc represents the combined proportion of alleles classified at the genus or better level (i.e., correct genus or correct species). For genomic context classification by CARD k-mers, categories include rejected (insufficient k-mers), unclassified, erroneous (incorrect genomic context), ambiguous (containing both chromosomal and plasmid k-mers), and correctly classified to a unique genomic context (chromosome or plasmid). Percentages represent the proportion of tested alleles in each category.

For genomic classification, CARD k-mers correctly identified the genomic origin (chromosome or plasmid) for 63.18% of chromosome and 20.02% of plasmid ARG alleles. A substantial proportion of alleles matched both genomic types (35.61% for chromosome and 77.47% for plasmid). Erroneous classification rates remained low (1.03% for chromosome, 2.5% for plasmid), and very few sequences were unclassified or rejected (Figure 4).

### Optimal k-mer Size

Pathogen classification accuracy plateaued at approximately 80% beyond a k-mer size of 15, with a sharp drop in unclassified predictions as k increased, while correct genus, erroneous, and rejected categories remained stable across larger k-mer sizes (Figure 5).

**Figure 5.**
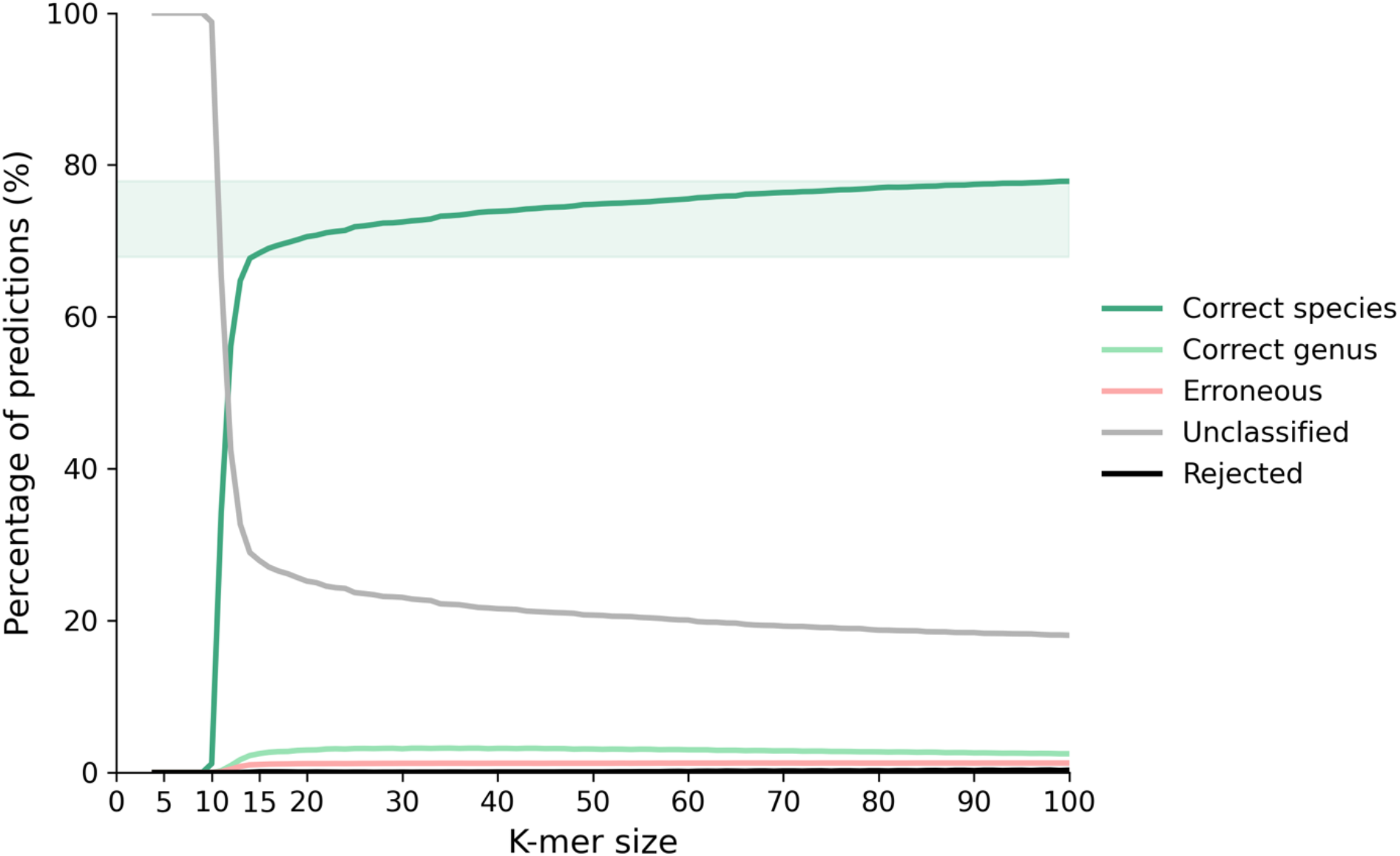
Effect of k-mer size on pathogen classification accuracy with CARD k-mers. Pathogen classification accuracy increases with k-mer size and plateaus around 80% beyond k=15. The proportion of unclassified predictions decreases notably as k increases, while correct genus, erroneous, and rejected categories remain relatively stable across k-mer sizes. The shaded region highlights accuracies within 10% of the maximum observed value, emphasizing the performance plateau and minimal gains in species-level accuracy beyond this range.

Genomic classification followed similar trends, and theoretical k-mer size analyses using entropy further supported a minimum k-mer size of 15 as a reliable minimum anchor size (Supplementary Figure 2, 3). Performance benchmarks show that unique 15-mers are built (*rgi kmer_build*) faster than 61-mers, with multi-threading reducing library construction time, but not memory usage. Query classification (*rgi kmer_query*) speed, measured in reads per minute, increased using 15-mers and multi-threading, while peak memory usage remained constant for each k-mer size regardless of thread count (Figure 6).

**Figure 6.**
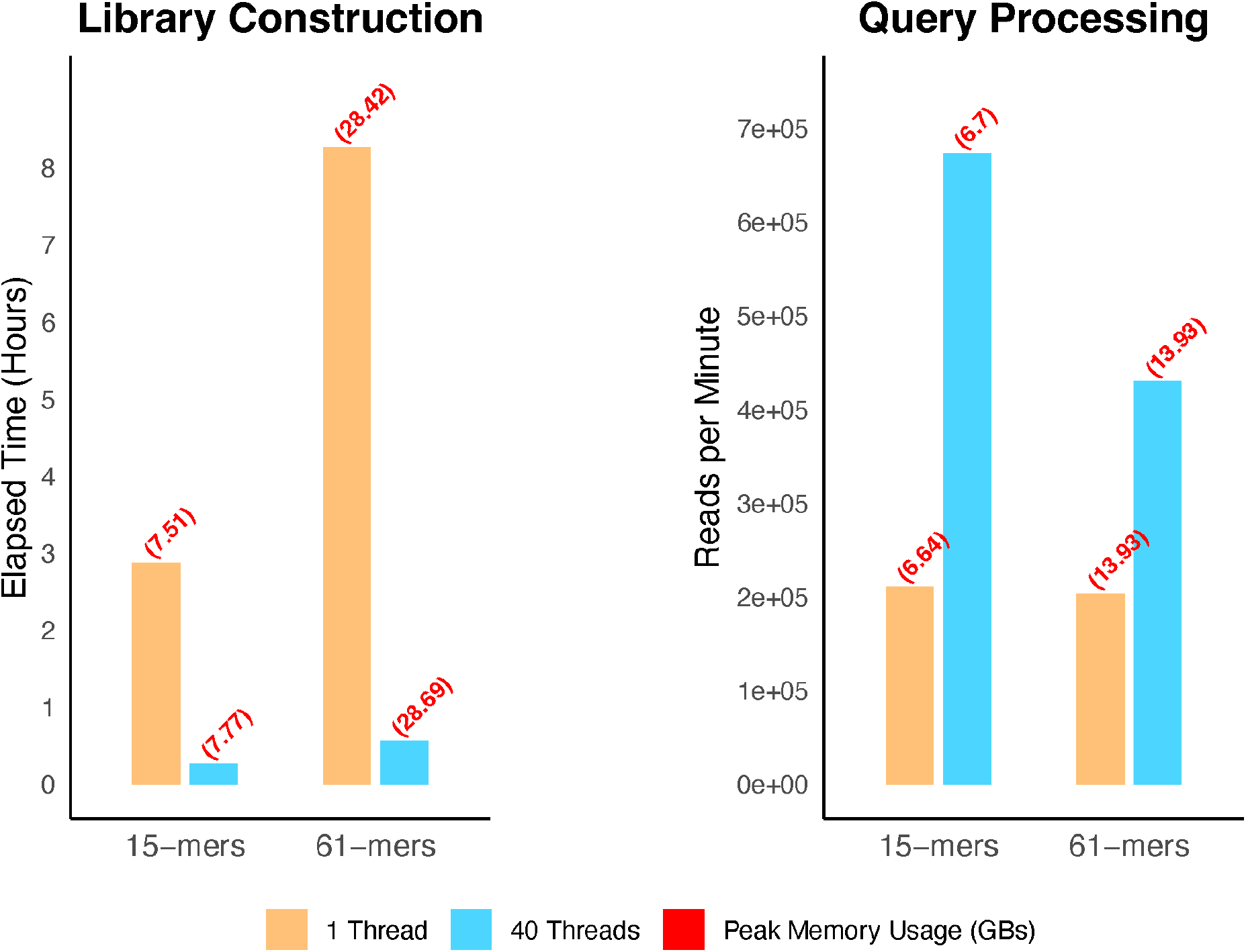
Performance benchmarks for CARD k-mers. K-mer library construction (left) and query processing (right) performance are shown for CARD k-mers using either 15-mers or 61-mers with single-thread (1 thread) or multi-threaded (40 threads) execution. Elapsed time (left) is measured in hours and represents total wall-clock time to build the reference library. Classification speed (right) is measured in reads per minute (RPM) for query operations. Peak memory usage, recorded as maximum resident set size (GB), is displayed as red text above each bar and was consistent across thread configurations for k-mer size.

## DISCUSSION

### Reliable Pathogen-of-Origin Prediction

CARD k-mers outperformed Kraken2 by 10.9% and CLARK by 15.2% in species accuracy for pathogen-specific AMR alleles, supporting its utility as an alignment-free tool for pathogen-of-origin prediction in sequences encoding ARGs. Although Kraken2 based on CARD reference sequences achieved 3.1% higher species accuracy than CARD k-mers, it produced 17.4% more erroneous calls. Erroneous calls may pose significant public health risks, as the misidentification of resistant pathogens can lead to inappropriate interventions (24). CARD k-mers adopts a conservative approach by tolerating a higher percentage of unclassified predictions to minimize false positives, ensuring a lower error rate than Kraken2, which aims to assign the maximum number of reads to species, resulting in a lower F1 score due to reduced assignment fidelity (25). The conservative paradigm of CARD k-mers is particularly valuable for AMR surveillance, where precision is paramount. If desired, CARD k-mers allows users to adjust parameters to balance sensitivity and specificity according to their surveillance needs, enhancing its versatility and applicability.

Additionally, CARD k-mers yielded the highest genus level accuracy at 3.0%. Genus-level AMR surveillance can play a useful role in guiding treatment and mitigation strategies, even when species-level identification is unavailable. Many bacterial genera exhibit characteristic resistance mechanisms that can inform clinical decision-making. For instance, *Pseudomonas* species possess intrinsic resistance to amoxicillin due to efflux pumps and porin modifications, which render the antibiotic ineffective (26). Early genus-level identification of *Pseudomonas* in clinical specimens allows clinicians to rule out the use of amoxicillin and other ineffective treatments, opting instead for targeted therapies such as piperacillin-tazobactam or ceftazidime while awaiting species-level data.

Further developments should focus on the validation of clinical and environmental metagenomic sequencing datasets (as opposed to whole alleles), and subsequent statistical analysis, such as the synergy implemented between Kraken2 and Bracken to estimate microbial abudance (27). This addition will enrich classifications performed by CARD k-mers, adding a quantitative dimension that deepens users’ understanding of the ARG landscape.

### Genomic Classification Contextualizes ARG Mobility Risk

The conservative paradigm of CARD k-mers additionally proved effective in reducing error rates during genomic classification. Genomic classification results in a high number of alleles assigned to the “plasmid & chromosome k-mers” category, a class we recommend retaining rather than binning them into the unclassified category. These k-mers, while not unique to one genomic context, still provide valuable mobility context and help prevent incorrect conclusions. For instance, even in the absence of direct plasmid k-mers, the presence of plasmid & chromosome k-mers precludes ruling out plasmid-borne AMR. Future updates to CARD k-mers will incorporate genomic islands, another type of mobile genetic element that harbors AMR genes (28) and that is available in the CARD-R reference data.

### Domain Specificity Improves Classification

Providing CARD-R AMR alleles as reference data to Kraken2 improved species prediction accuracy by 14%, reduced the number of erroneous calls by 13.1%, and reduced the number of unclassified calls by 1.5%, despite the standard Kraken2 database being over 70GB larger than CARD-R. This finding suggests that k-mer classifier accuracy can be significantly improved by defining a precise sequence space to build the required reference k-mers, and extends previous work done on Kraken2 general database optimization to the AMR sequence space (18, 19).

### Recommended k-mer Size

K-mer size is an arbitrary yet important parameter for k-mer classifiers (29). As a default, CARD k-mers uses 61-mers, Kraken2 35-mers, CLARK 31-mers, but there is limited empirical evidence to support these choices. Based on our CRE curves, experimental data and performance monitoring, we recommend either 15-mers for faster library building and query processing, or 61-mers for a slightly higher accuracy setting, albeit at worse computational performance. These findings align with recent works investigating k-mer based ARG detection, which suggest smaller sized 13-mers (30). We do not recommend using sizes smaller than 15-mers, and sizes that are too large risk being larger than the ARGs themselves, resulting in those sequences being rejected. Furthermore, the effects of smaller k-mers on variant AMR sequences and their subsequent classification are largely unknown and require further investigation; it is plausible that smaller k-mers offset the effects of single nucleotide variants (SNVs) since k-mer coverage can instead occur over conserved regions of the AMR alleles. As of September 2025, the CARD-R data set includes both pre-compiled 15-mer and 61-mer reference libraries.

### CARD k-mers Performance

Overall, CARD k-mers sacrifices speed for improved classification accuracy compared to Kraken2 and CLARK in terms of reads per minute, but due to its highly specialized library, achieves k-mer library construction much faster than Kraken2 and CLARK, which can take several days to build. CARD will shoulder the burden of library construction by continuing to provide pre-compiled k-mer libraries. Additionally, users can pre-process their input data to contain only ARGs by using RGI before classifying them with CARD k-mers, and take advantage of multi-threading if possible. The eventual implementation of faster k-mer counting algorithms into CARD k-mers will improve speed and reduce memory usage (31).

### Mobility and Evolution of AMR Alleles

The LGT of ARGs among bacteria complicates the accurate taxonomic classification of short metagenomic reads. Gene mobility enables diverse bacterial species to acquire and share ARGs, leading to discrepancies between phylogenetic relationships and the distribution of resistance phenotypes. Consequently, metagenomic analyses may misattribute ARGs to incorrect taxa, posing challenges in identifying the true reservoirs and vectors of AMR. Understanding the extent and mechanisms of LGT is crucial for improving the precision of metagenomic AMR surveillance and developing effective strategies to combat the spread of resistance (32).

CARD k-mers addresses some of the challenges introduced by LGT by leveraging its pathogen-centric AMR allele database (414 pathogens as of CARD-R 4.0.2), which stores only unique pathogen k-mers from CARD-R. This approach inherently includes SNVs for AMR alleles associated with distinct pathogens. These embedded pathogen-specific SNVs act as genetic markers, enabling the differentiation of native AMR alleles from those acquired via LGT, partially mitigating concerns about gene mobility (33, 34). Additionally, 310,891 (97%) of the AMR alleles in CARD-R have a single host pathogen, enabling precise taxonomic predictions for most alleles (Supplementary Figure 4). For the remaining 3% of mobile AMR alleles, users can still assess genomic context and flag these ARGs for further analysis. SNVs also present significant challenges in the classification of emerging AMR alleles.

The presence of SNVs can lead to variations that are difficult to interpret, complicating the determination of an allele’s functional impact. This complexity is further exacerbated by the limitations of current classification methods, which may not adequately account for the diverse effects of SNVs on gene function (35). However, CARD k-mers may better tolerate SNVs over other classifiers via the construction and storage of ARG variant-specific k-mers. This capability is enabled by the inclusion of “Strict” alleles in CARD-R, which represent ARG variants not yet been described in the literature. As a result, CARD k-mers is uniquely positioned to detect both known and emerging ARGs in pathogens, making it a powerful tool for comprehensive AMR surveillance.

Future iterations of CARD k-mers have the potential to provide rich phenotypic insights by integrating the extensive annotations of ARGs available in CARD. This will enable deeper k-mer analyses, such as predicting affected drug classes and resistance mechanisms. These enhancements could improve the genotype-to-phenotype predictive power of CARD k-mers for short metagenomic reads.

### Classifying Unclassifiable Alleles

Despite the strategies CARD k-mers implements to mitigate LGT, not all AMR alleles receive classification. UMAP analysis of the 20,456 unclassified AMR alleles revealed two main clusters associated with *E. coli* and *K. pneumoniae*, driven by shared resistance profiles and conserved genes like *Escherichia coli* EF-Tu mutants conferring resistance to pulvomycin (*Ecol_EFTU_PLV)*, sulfonamide resistant dihydropteroate synthase *sul1*, and tetracycline efflux pump *tet(A)* (Supplementary Figure 5, Table 2). *EF-Tu*, a conserved bacterial housekeeping gene encoding elongation factor Tu, is difficult to classify due to shared k-mers across bacterial species (36). Similarly, *sul1*, a highly mobile antibiotic resistance gene that has propagated globally since the 1940s, reflects the selective pressures exerted by the widespread use of early synthetic antimicrobials, such as sulfonamides. Its extensive mobility and wide pathogen distribution complicate efforts to predict its pathogen-of-origin, thereby challenging the tracing of its evolutionary and epidemiological history (37).

These findings underscore the inherent difficulty in classifying conserved genes and mobile ARGs with redundant k-mers. They highlight the limitations of current classification strategies and emphasize the need for additional contextual data, such as genomic location, association with mobile genetic elements (MGEs), and phylogenetic analysis to enhance classification accuracy. Future efforts should integrate complementary epidemiological methods to refine the identification of AMR alleles, particularly for those that remain ambiguous due to their high mobility or shared evolutionary history.

## CONCLUSIONS

Overall, CARD k-mers represents a much-needed advancement over current methods in ARG surveillance by addressing key challenges in pathogen-of-origin identification and ARG mobility assessment. Leveraging the pathogen-specific CARD-R database, it achieves high taxonomic specificity and detects pathogens for both known and novel ARG variants through the inclusion of “Strict” allele annotations in CARD-R, offsetting the effects of LGT and SNVs on classification. Additionally, its genomic context analysis via plasmid- and chromosome-specific k-mers provides valuable insights into the dissemination potential of ARGs. Further refinements to the algorithm and streamlined workflows should be investigated to improve classification accuracy and expand genotype-to-phenotype predictions, ultimately strengthening untargeted AMR surveillance and improving clinical outcomes.

## SOFTWARE & DATA AVAILABILITY

Version controlled copies of the Resistance Gene Identifier (RGI) are available at the CARD GitHub repository: https://github.com/arpcard/rgi. All data files for validation can be found in GitHub repository https://github.com/mawlodarski/card-kmers. Benchmarking data be found at https://zenodo.org/records/6543357.

## SUPPLEMENTARY DATA

Supplementary Data are available in association with this manuscript at *bioRxiv*.

## FUNDING

This study was supported by the Canadian Institutes of Health Research (PJT-156214 to AGM), Genome Canada (to RGB), and funds from the Comprehensive Antibiotic Resistance Database. TTYL was supported by a Michael G. DeGroote Institute for Infectious Disease Research Summer Student Fellowship. TET was supported by an Ontario Graduate Fellowship. FM was supported by a Donald Hill Family Fellowship in Computer Science.

## ACKNOWLEDGEMENTS

Computational support was provided by the McMaster Faculty of Health Sciences Advanced Computing Facility, supplemented by hardware donations and loans from Cisco Systems Canada, Hewlett Packard Enterprise, and Pure Storage. AGM holds McMaster’s David Braley Chair in Computational Biology, generously supported by the family of the late Mr. David Braley. Thank you to Drs. Samantha L. Wilson and Jonathan M. Stokes for their guidance and mentorship of this work.

## CONFLICT OF INTEREST

The authors declare that there are no conflicts of interest.

## AUTHOR CONTRIBUTIONS

MAW: conceptualization, methodology, investigation, software, data curation, formal analysis, visualization, writing – original draft, writing – review and editing. TTYL: conceptualization, methodology, software, data curation, visualization, writing – review and editing. BPA: data curation, supervision, writing – review and editing. ARR: software, supervision, writing – review and editing. TET: visualization, writing – review and editing. FM: conceptualization, methodology, writing – review and editing. RGB: conceptualization, funding acquisition, project administration, writing – review and editing. AGM: conceptualization, funding acquisition, project administration, supervision, writing – review and editing.

## Notes

https://github.com/arpcard/rgi

https://zenodo.org/records/6543357

https://github.com/mawlodarski/card-kmers

### Competing Interest Statement

The authors have declared no competing interest.

