## Supplementary Figures & Tables for "CARD k-mers: Unmasking the pathogen hosts and genomic contexts of antimicrobial resistance genes in metagenomic sequences"

### SUPPLEMENTARY FIGURES AND TABLES

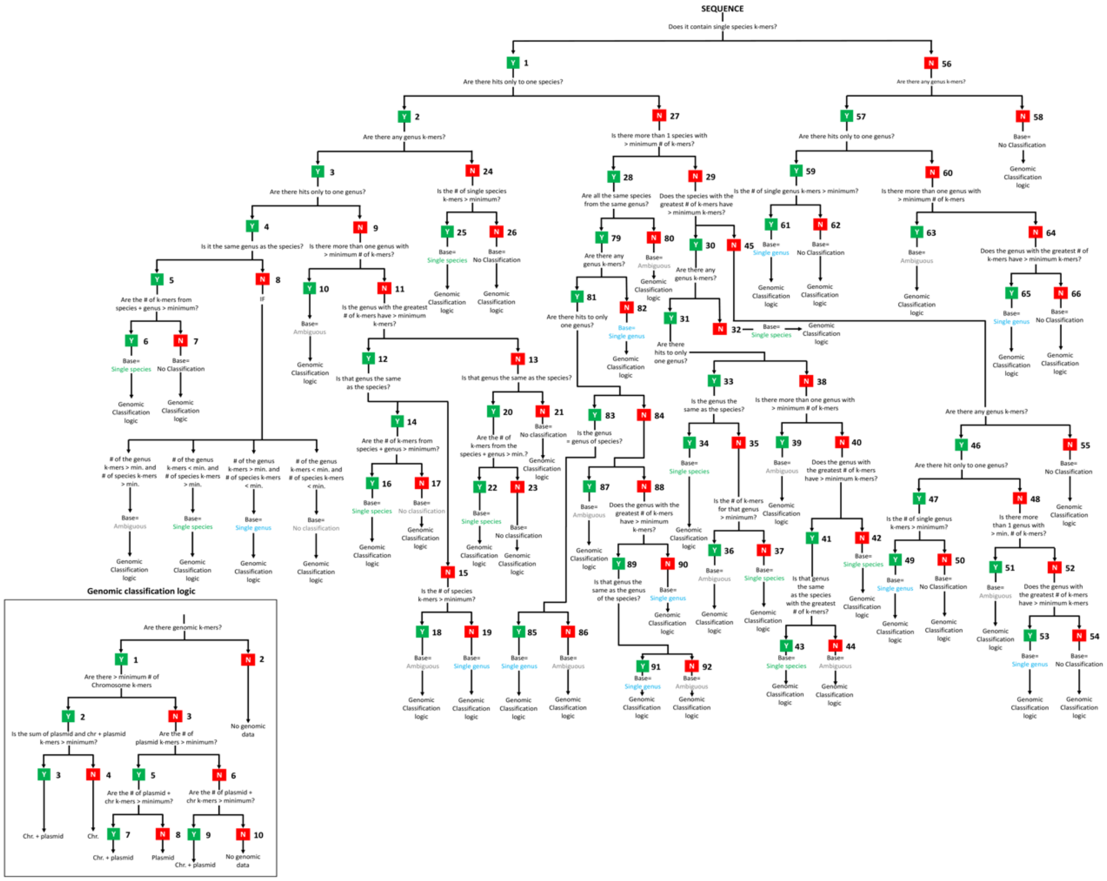


**Supplementary Figure 1: CARD k-mers classification logic tree.** To receive pathogen and genomic classification, input sequences are passed through this rules-based decision tree. The first goal is to provide the query sequence with a "base" value. Four possible base values exist: single species, single genus, ambiguous and no classification. Once a base value has been assigned, CARD k-mers predicts the ARG's genomic location. Four possible genomic predictions exist: chromosome, plasmid, chromosome and plasmid, or no genomic data. A more high-definition version can be visualized at https://github.com/arpcard/rgi.

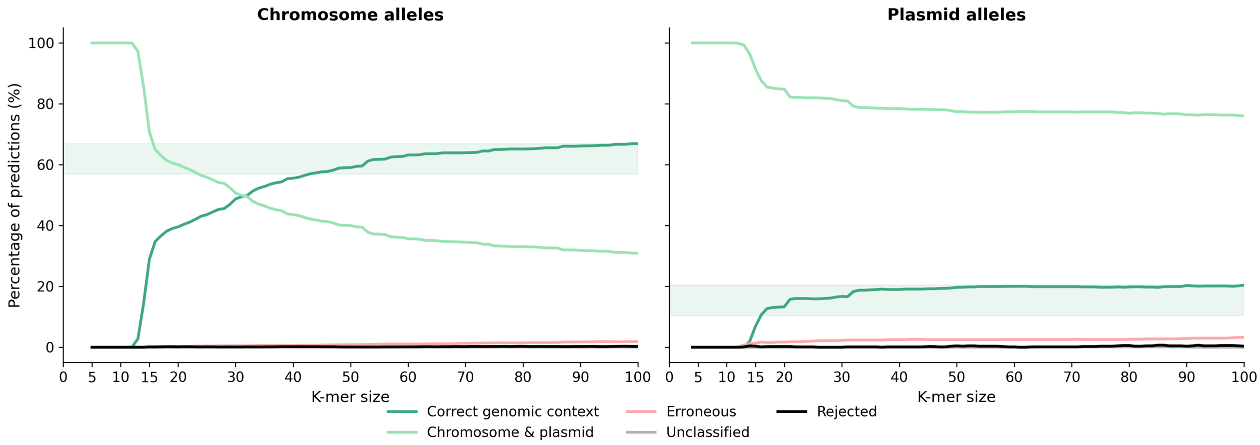


**Supplementary Figure 2. Effect of k-mer size on genomic context classification accuracy with CARD k-mers.** Genomic classification performance is shown for chromosome alleles (left) and plasmid alleles (right). For both, the proportion of correct genomic context predictions increases with k-mer size and eventually plateaus. Sequences containing both chromosome and plasmid k-mers decrease as correct predictions rise, highlighting an inverse relationship between ambiguous and definitive context calls. The shaded regions represent accuracy values within 10% of the maximum correct genomic context rate, indicating where further increases in k-mer size yield diminishing improvements. Other prediction categories (erroneous, unclassified, and rejected) remain low and stable across all k-mer sizes.

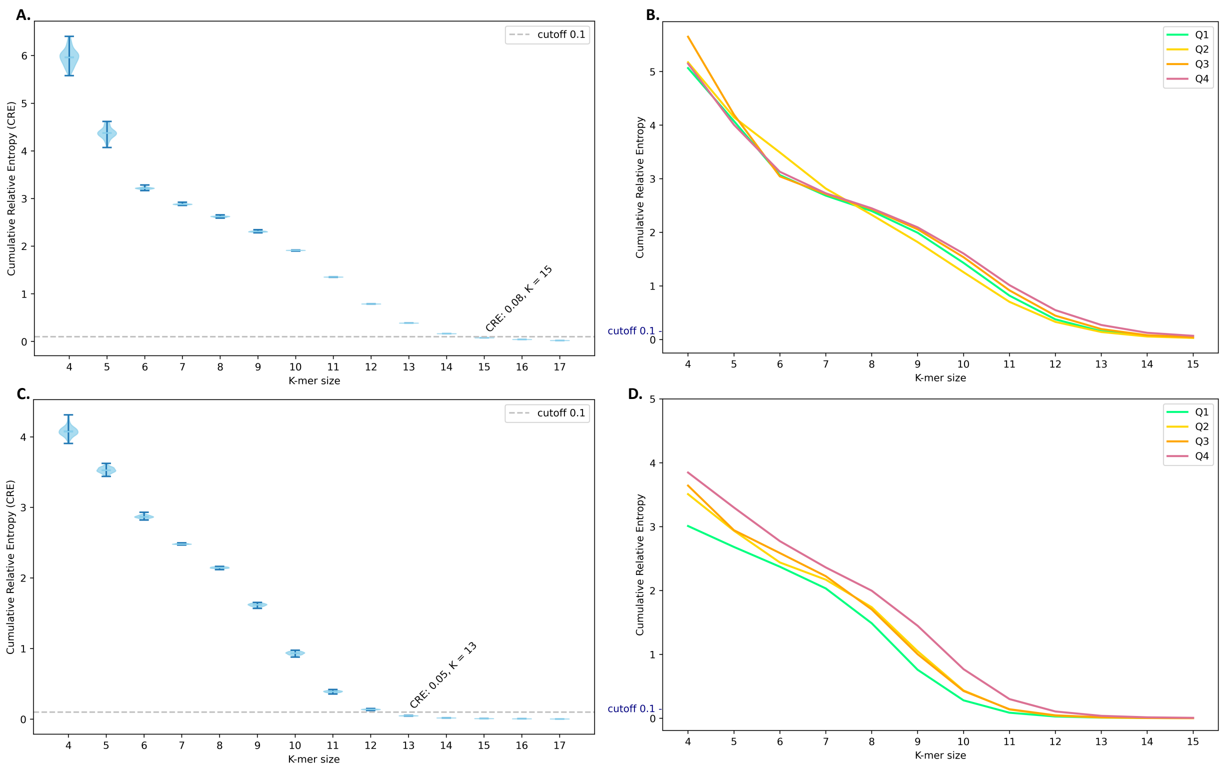


**Supplementary Figure 3: Cumulative relative entropy (CRE) analysis of CARD-R AMR alleles:** 103,456 AMR alleles from the CARD-R testing set were stratified into size quartiles (Q1: smallest – Q4: largest) and bootstrapped into 100 sub-datasets, totaling 10,338,900 *in silico* alleles. Using KITSUNE v1.3.1, the average CRE for pathogen-specific alleles was calculated in (A), and the CRE for each quartile was plotted in (B). The same analysis was performed for 89,300 plasmid-specific AMR alleles, with results shown in (C) and (D). A CRE cutoff value of 0.1 was used to determine the optimal minimum k-mer size.

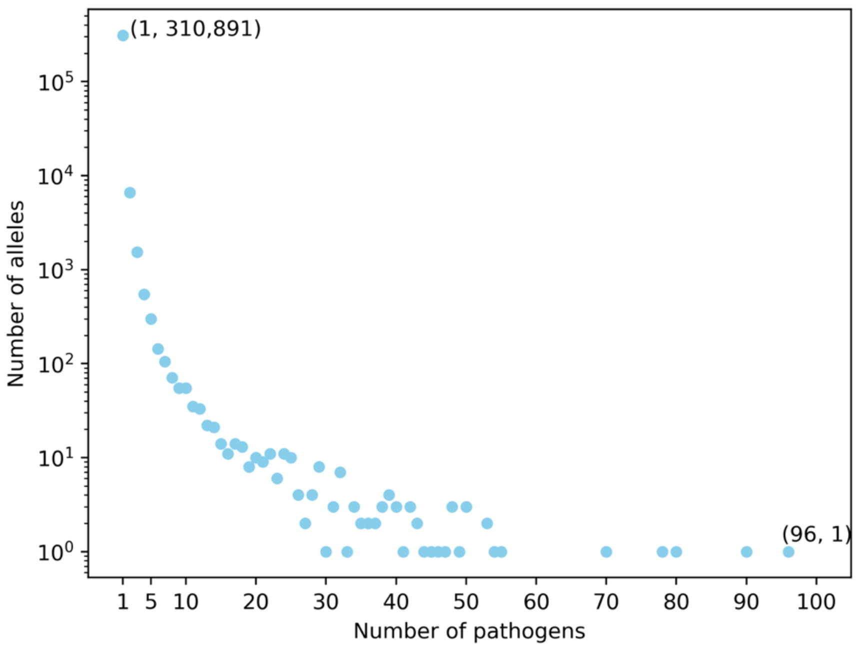


**Supplementary Figure 4: Distribution of CARD-R v4.0.0 alleles**. The graph illustrates the number of distinct pathogens encoding each unique allele. The y-axis (logarithmic scale) represents the number of unique alleles, and the x-axis denotes the number of pathogens encoding each allele count. Of the total alleles in CARD-R, 310,891 were found in only one species, while 9,722 were identified in more than one species. One notable allele, *adeF*, a membrane fusion protein of the multidrug efflux complex AdeFGH, was found in 96 distinct pathogens. Widely distributed AMR alleles are better suited for genomic classification with CARD k-mers, as opposed to pathogen classification.

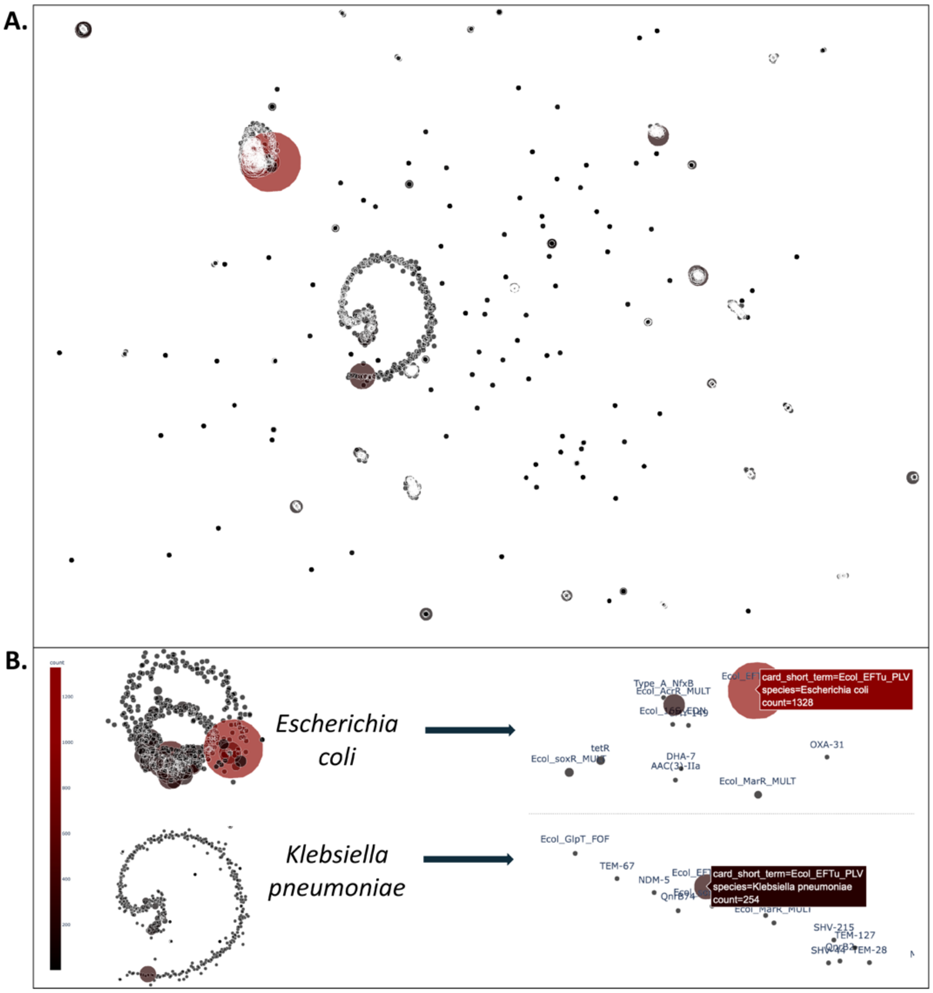


**Supplementary Figure 5. UMAP visualization of unclassified AMR alleles from CARD k-mer classification validation.** (A) A total of 20,462 AMR alleles labeled as unclassified during pathogen classification were vectorized into a binary allele-pathogen matrix, where each allele was represented by its association across known CARD-R pathogens. This matrix was used for dimensionality reduction via UMAP. (B) The resulting 3D UMAP projection revealed two major clusters corresponding to *Escherichia coli* and *Klebsiella pneumoniae*-associated alleles. The most abundant allele in both clusters was *Ecol_EFTU_PLV*, a conserved housekeeping gene encoding elongation factor Tu (EFTu), found 1,328 times in *E. coli* and 254 times in *K. pneumoniae*. Full interactive UMAP visualization available at: https://github.com/mawlodarski/card-kmers.

| Code | K-mer Type | Count |
|---|---|---|
| **P** | Plasmid | 295,432 |
| **C** | Chromosome | 12,225,582 |
| **B** | Plasmid + Chromosome | 603,319 |
| **S** | Species | 27,401,594 |
| **G** | Genus | 1,259,938 |
| | | **40,525,927** |

**Supplementary Table 1: CARD k-mers reference library k-mer counts.** Over 40 million 61-mers are built and stored for taxonomic and genomic predictions. P, C, B class k-mers are used to predict the genomic location of in input ARG sequence data, while S and G class k-mers are used to predict pathogen-of-origin.

| CARD AMR Gene | Resistance Mechanism | # of Pathogens in CARD-R with Gene |
|---|---|---|
| ***Ecol_EFTU_PLV*** | Target alteration | 87 |
| ***evgS*** | Efflux | 24 |
| ***sul1*** | Target replacement | 95 |
| ***tet(A)*** | Efflux | 61 |
| ***tet(M)*** | Target protection | 100 |

**Supplementary Table 2: List of the 5 most abundant unclassified AMR alleles by CARD k-mers.** Both mobile resistance genes (e.g., *sul1, tet(A), tet(M))* and conserved housekeeping genes (e.g., *EFTu*) pose classification challenges due to their widespread distribution and sequence similarity across bacterial species.
